# Missense variants causing Wiedemann-Steiner syndrome preferentially occur in the KMT2A-CXXC domain and are accurately classified using AlphaFold2

**DOI:** 10.1101/2022.01.03.474770

**Authors:** Tinna Reynisdottir, Kimberley Anderson, Leandros Boukas, Hans Tomas Bjornsson

## Abstract

Wiedemann-Steiner syndrome (WSS) is a neurodevelopmental disorder caused by *de novo* variants in *KMT2A*, which encodes a multi-domain histone methyltransferase. To gain insight into the currently unknown pathogenesis of WSS, we examined the spatial distribution of likely WSS-causing variants across the 15 different domains of KMT2A. Compared to variants in healthy controls, WSS variants exhibit a 64.1-fold overrepresentation within the CXXC domain – which mediates binding to unmethylated CpGs – suggesting a major role for this domain in mediating the phenotype. In contrast, we find no significant overrepresentation within the catalytic SET domain. Corroborating these results, we find that hippocampal neurons from *Kmt2a*-deficient mice demonstrate disrupted H3K4me1 preferentially at CpG-rich regions, but this has no systematic impact on gene expression. Motivated by these results, we combine accurate prediction of the CXXC domain structure by AlphaFold2 with prior biological knowledge to develop a classification scheme for missense variants in the CXXC domain. Our classifier achieved 96.0% positive and 92.3% negative predictive value on a hold-out test set. This classification performance enabled us to subsequently perform an *in silico* saturation mutagenesis and classify a total of 445 variants according to their functional effects. Our results yield a novel insight into the mechanistic basis of WSS and provide an example of how AlphaFold2 can contribute to the *in silico* characterization of variant effects with very high accuracy, establishing a paradigm potentially applicable to many other Mendelian disorders.

## Introduction

Wiedemann Steiner syndrome (WSS, OMIM: 605130) is a Mendelian disorder of the epigenetic machinery. Its phenotypic features include intellectual disability, postnatal growth deficiency, hypertrichosis, and characteristic facial features. WSS is typically caused by heterozygous *de novo* variants in the gene encoding the histone-lysine N-methyltransferase 2A (KMT2A) [1]. *KMT2A*, also known as *MLL*/*MLL1*, is post-translationally cleaved into N-terminal and a C-terminal fragments, which subsequently heterodimerize. Each of these fragments contains several annotated protein domains. The larger N-terminal fragment contains three AT-hooks, a cysteine-rich CXXC domain, four plant homeodomain (PHD) fingers, a bromo domain and a FYR-N domain. The smaller C-terminal fragment contains a transactivation (TAD) domain, a *Win* motif, FYR-C domain, a SET, and a post-SET domain [2].

In this study, we address two questions. First, how important is the role of the different KMT2A domains in the pathogenesis of WSS? The answer would provide important clues into the molecular basis of the disorder, since the different domains mediate different functions. Second, what are the rules that determine how the genetic disruption of the most important domain(s) causes WSS? The answer would enable a systematic characterization of WSS-causing variants, informing basic biology as well as future clinical decision-making.

To answer the first question, we adopt an unbiased genetic approach. We examine the spatial distribution of likely WSS-causing missense variants across the different domains, reasoning that such variants will be enriched in the domains most critical for WSS pathogenesis when compared against variants in healthy individuals. To address the second question, we focus on the CXXC domain responsible for binding unmethylated CpGs [3], which we find shows by far the greatest enrichment. We combine accurate structure prediction by AlphaFold2 with existing biological data to create an effect classification scheme for missense variants in the CXXC domain. After validating our classifier, we deploy it to perform an *in silico* saturation mutagenesis, examining a total of 445 variants.

## Results

### Preferential occurrence of missense variants likely pathogenic for Wiedemann-Steiner syndrome in the CXXC domain of KMT2A

If certain domains of a protein are critical for its function, disease-causing missense variants will tend to preferentially occur in these domains. We thus set out to explore the distribution of *KMT2A* missense variants (MVs) across its 15 different domains. We started by identifying 57 MVs that are likely pathogenic for WSS (**Methods; Figure 1A**). As a control, we used 1340 MVs which are present in gnomAD and are therefore not expected to cause WSS (**Methods; Figure 1A**). We tested each KMT2A domain for enrichment of WSS MVs relative to gnomAD MVs. We discovered a 64.1-fold enrichment in the CXXC domain (**Figure 1B**; Fisher’s exact test, Bonferroni-adjusted p = 5.5e-17). This far exceeded the enrichment observed at other domains, with the second most enriched domain being the fourth PHD finger (**Figure 1B**; 18.4-fold enrichment; Fisher’s exact test, Bonferroni-adjusted p = 0.03), and the first PHD finger also showing significant enrichment (**Figure 1B**; 8.5-fold enrichment; Fisher’s exact test, Bonferroni-adjusted p = 0.01).

**Figure 1.**
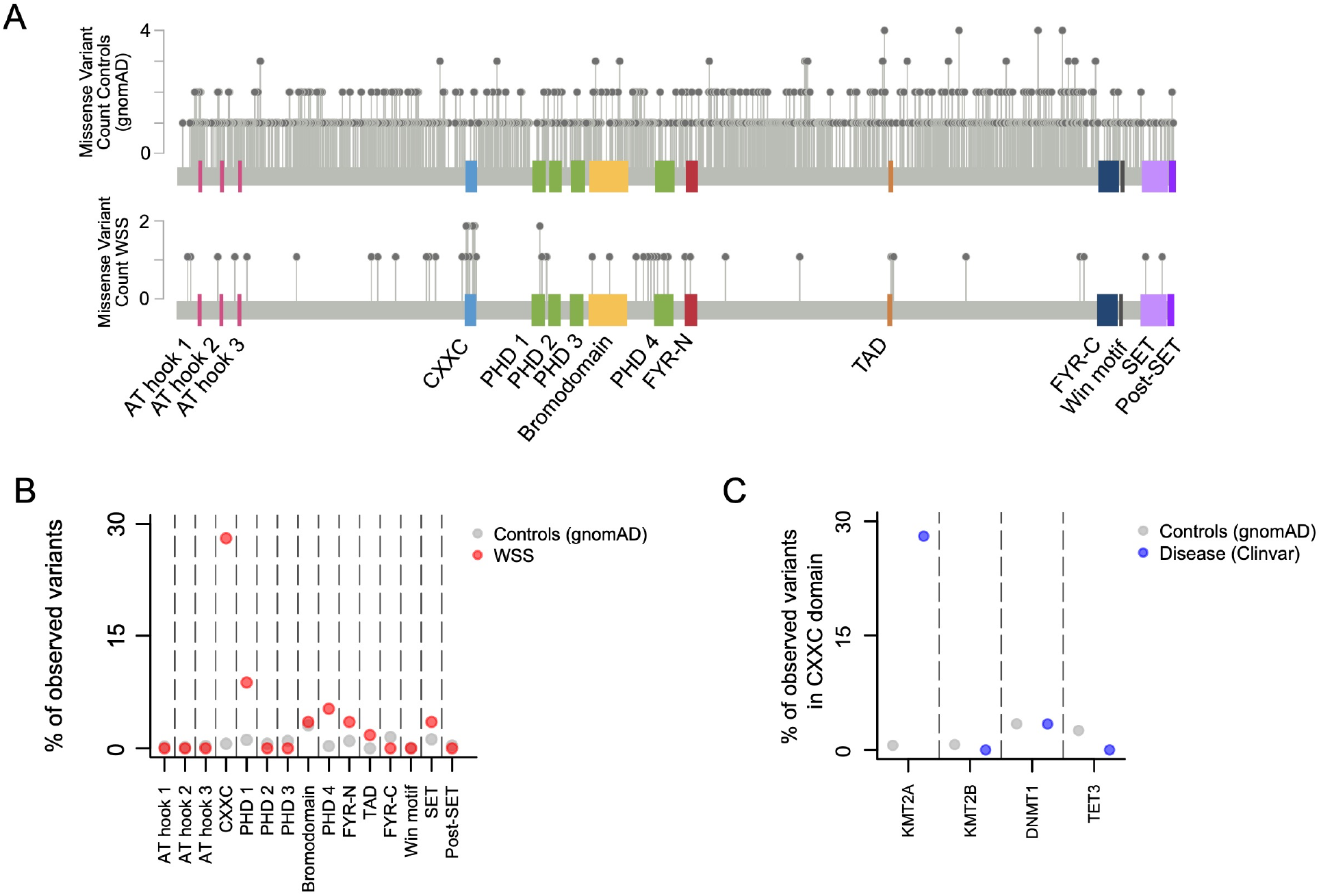
The distribution of likely pathogenic Wiedemann-Steiner syndrome missense variants across the different domains of KMT2A. **(A)** KMT2A missense variants in gnomAD (top) and WSS (bottom). See Methods for filtering criteria. **(B)** The percentage of missense variants from gnomAD (grey dots) and WSS (red dots) that fall in each of the different domains of KMT2A. **(C)** The percentage of missense variants in gnomAD (grey dots) and variants with evidence of pathogenicity (blue dots) that fall in the CXXC domain of different epigenetic regulators.

To assess the robustness of our result, we repeated our analysis using a control set of 1788 *KMT2A* somatic variants obtained from sequencing of tumor samples (**Methods**). While these variants are likely a mix of driver and passenger variants, the latter are expected to comprise the majority. We recapitulated our result, with the rank ordering of the different domains with respect to their enrichment of WSS MVs remaining unchanged (**Supplemental Figure 1A, B, C**). Across all domains, the enrichment estimates were attenuated compared to the gnomAD-vs-WSS comparison (**Supplemental Figure 1C)**. This is consistent with the pool of somatic variants containing a minority of cancer drivers, and also suggests that the same domains that contribute to WSS pathogenesis contribute to the tumorigenic role of KMT2A as well.

Notably, the catalytic SET domain of *KMT2A* does not show significant enrichment for WSS variants (**Figure 1B**; Fisher’s exact test, p = 0.165). In contrast, it shows significant, albeit weak, enrichment, when the somatic cancer variants are compared against the gnomAD controls (odds ratio = 2.87, Fisher’s exact test, p = 8.43e-05).

### The CXXC domains of other epigenetic regulators do not show enrichment for disease-causing missense variants

We next asked if the preferential occurrence of disease-causing missense variants in the CXXC domain is unique to *KMT2A*, or whether this is a general phenomenon across epigenetic regulators. Apart from *KMT2A*, three other CXXC-domain-containing epigenetic regulators have been linked to Mendelian diseases: *KMT2B* (DYS28; OMIM:617284), *DNMT1* (HSN1E; OMIM:614116), and *TET3* (BEFAHRS; OMIM:618798). However, we found that none of these genes exhibits significant enrichment of disease-causing MVs in the CXXC domain (**Figure 1C**; Methods; Fisher’s exact test, *DNMT1* p = 1, *KMT2B* p = 1, *TET3* p = 1), and verified that this is not due to inadequate power (**Methods**).

### The disruption of H3K4me1 in *Kmt2a*-deficient mice preferentially occurs at CpG-rich regions but has no systematic effect on gene expression

Since the CXXC domain mediates binding to clusters of unmethylated CpG dinucleotides [3], our domain enrichment results imply that KMT2A exerts its most important function at CpG-rich regions. Our results also imply that its most important function is not its catalytic activity as a histone methyltransferase, given the lack of MV enrichment in the SET domain. We sought to test these implications, by comparing genome-wide H3K4me1 and gene expression patterns in hippocampal CA (Cornu Ammonis) neurons from mice with *Kmt2a* knockout in excitatory neurons (*Kmt2a* cKO) and wild-type mice (using ChIP-seq and RNA-seq data; **Methods**) [4].

First, we found a strong relationship between the CpG-richness of a region and the probability of H3K4me1 disruption; CpG-rich peaks are much more likely to exhibit disruption upon *Kmt2a* cKO (5 wild-type vs 3 cKO mice; p<2.2e-16; **Figure 2A, B**; **Methods**). However, although regions with the strongest evidence for H3K4me1 disruption (1^st^ p-value decile) show a 34.4-fold enrichment at promoters (within 1 kb of the transcription start site on either side) compared to regions with the lowest evidence (10^th^ p-value decile; Fisher’s exact test, p<2.2e-16; **Supplemental Figure 2A**), we found no evidence for a systematic impact of H3K4me1 disruption on gene expression (5 wild-type vs 6 cKO mice; **Methods**). Specifically, genes whose promoters have disrupted H3K4me1 are not significantly more likely to be differentially expressed compared to genes without H3K4me1 disruption at their promoter (p = 0.47 for a shift in p-value distribution between the 1^st^ and 10^th^ deciles; **Figure 2C, D**; **Methods**). Taken together, these results show that: a) KMT2A preferentially acts at high-CpG-density regions, and b) its catalytic activity has little effect on gene expression, suggesting that it may also not affect higher-level phenotypes.

**Figure 2.**
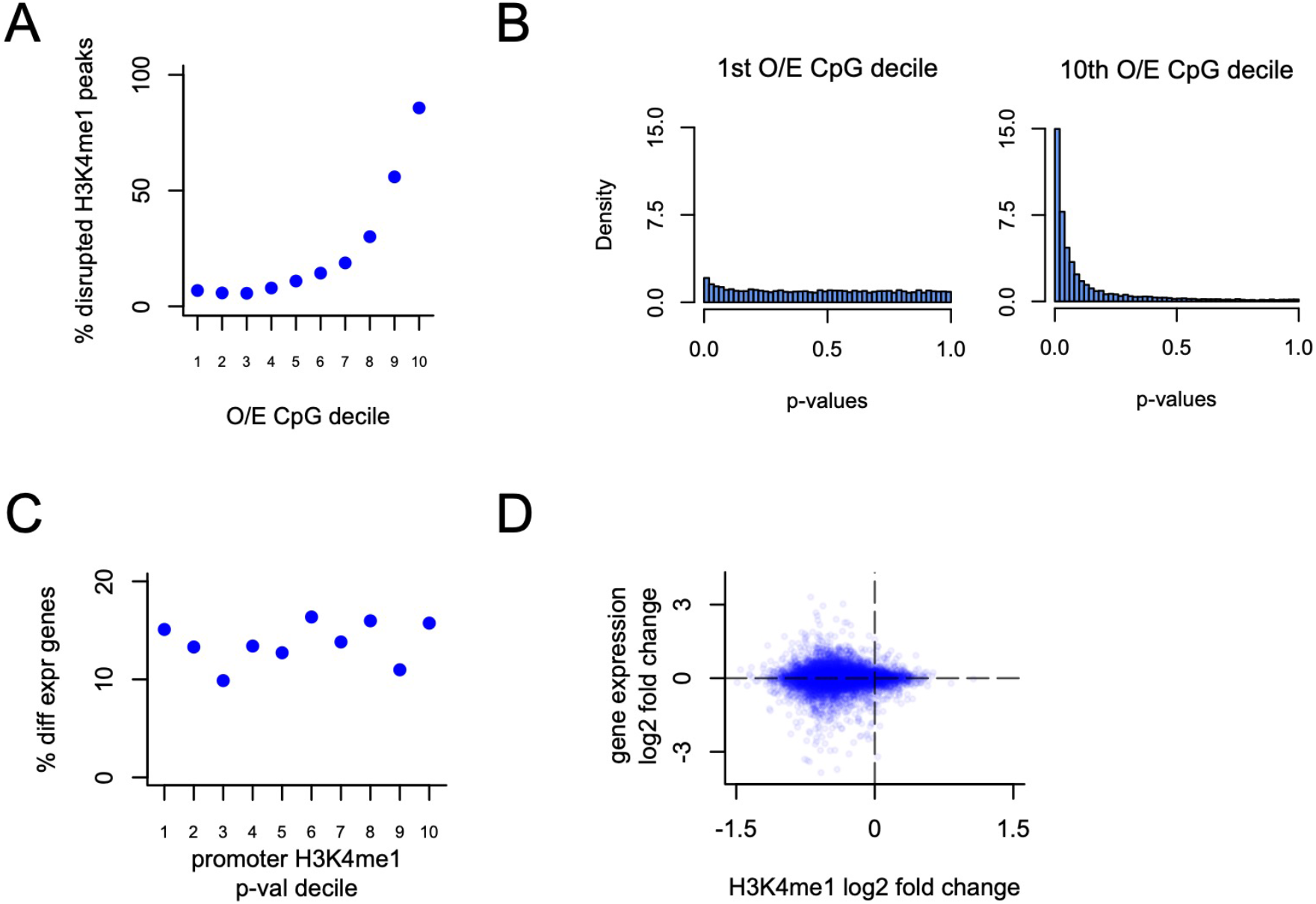
The relationship between disrupted H3K4me1, regional observed-to-expected CpG ratio, and gene expression in *Kmt2a* deficient mice. **(A)** The percentage of disrupted H3K4me1 peaks, stratified based on the observed-to-expected CpG ratio of the underlying peak sequence. **(B)** The p-value distributions from the H3K4me1 differential analysis, for the top and bottom observed-to-expected CpG ratio deciles of the peak sequences. **(C)** The percentage of differentially expressed genes, stratified based on the p-value of associated promoter peaks (+/- 1kb from the TSS) from the differential H3K4me1 analysis. **(D)** Scatterplot of the log2 fold change of H3K4me1 peaks against the log2 fold change of downstream gene expression. Each point corresponds to a gene-promoter pair. In cases where multiple peaks were present at the same promoter, the average log2 fold change was computed.

### An AlphaFold2-based scheme classifies missense variants in the CXXC domain of KMT2A with high accuracy

Given our evidence for a central role of the KMT2A-CXXC domain in WSS, we sought to develop a variant classification scheme that would: a) label any possible variant in the CXXC domain as pathogenic or not, and b) in the case of pathogenic variants, provide a characterization of their functional effect. We reasoned that such a classifier should take into account the effect of variants on the 3-dimensional structure of the domain, the mean Coulombic electrostatic potential of the mutant structure, and the ability of the mutated domain to form hydrogen bonds with the DNA backbone.

To assess the feasibility of our approach, we first examined if AlphaFold2 (AF2) - which has recently enabled the determination of 3D-protein structures with experimental-level accuracy - accurately predicts the structure of the CXXC domain. We observed a highly confident prediction (**Figure 3A**; pLDDT>70 for 96.5% of residues and pLDDT>90 for 82.5% of residues). In addition, all features previously identified in solution and crystal structures of the domain (ID: 2j2s, 2jyi, 4nw3, 2kkf) were present in the predicted structure: a crescent overall shape, two antiparallel beta sheets at the N and C terminals, and four alpha helices (**Figure 3B**). While no single experimentally derived structure contains all these features, this can be explained by the domain existing in multiple conformational states.

**Figure 3.**
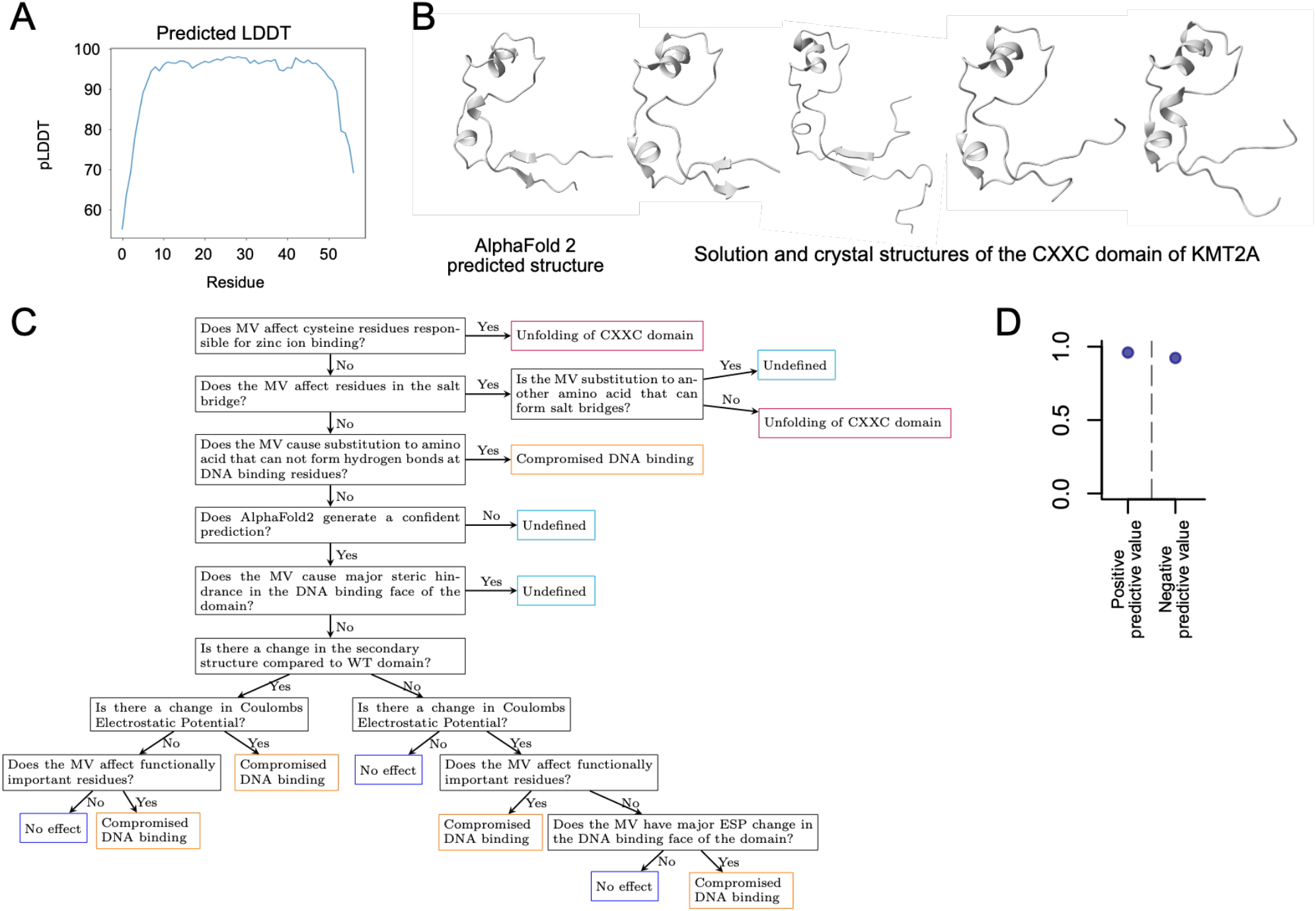
An AlphaFold2-based variant effect classification scheme for the CXXC domain of KMT2A. **(A)** Predicted LDDT values for the CXXC domain of KMT2A. **(B)** The AlphaFold2-predicted and experimentally determined structures of the CXXC domain of KMT2A. **(C)** The variant effect classification scheme. Functionally important residues are defined as residues that have been experimentally determined to be implicated in electrostatic interaction with the DNA backbone, residues responsible for hydrogen bond formation with the DNA and residues at the structurally important KFGG site. **(D)** The positive and negative prediction value that the classifier shown in **(C)** attained on a hold-out test set consisting of 38 missense variants (see text for details).

We then proceeded to derive a classification scheme, using a training set of 14 MVs with experimentally determined effects and prior biological knowledge about the domain (**Figure 3C**; **Supplemental Figure 4A, Supplemental Table 3**) [5]. The first steps of the scheme pertain to residues whose substitution effects are not captured by AF2 (**Methods**): the eight cysteines responsible for zinc ion binding, the residues forming the salt bridge, and the residues that form direct hydrogen bonds with the DNA. The rest of the scheme is then divided into two cascades, based on the AF2-predicted mutant secondary structure and its mean Coulombic electrostatic potential (**Methods**). Variants receive different classifications depending on their effect on secondary structure/electrostatic potential, as well as their position.

To evaluate the performance of our classifier, we used a hold-out test set consisting of 38 MVs. Of these, 16 were MVs with strong evidence for pathogenicity for WSS (**Methods**), 13 were MVs with experimentally determined effects [5, 6], and 9 were MVs seen in gnomAD/TOPMed, and are thus expected to be benign (**Supplemental Figure 4B, Supplemental Table 3**). On this test set, our classifier attained a 96.0% positive and a 92.3% negative prediction value (**Figure 3D**). Notably, out of the 16 WSS MVs present in the CXXC domain of *KMT2A*, nine are positioned at cysteine residues responsible for zinc ion binding compared to none of the control variants (Fisher’s exact test, p = 0.009496).

### *In silico* saturation mutagenesis classifies 445 variants in the CXXC domain of KMT2A

The accuracy with which our classifier performs motivated us to perform an *in silico* saturation mutagenesis for the CXXC domain of KMT2A, with the goal of creating a resource that would enable rapid classification of newly encountered variants in patients. We focused on the 50/57 residues that have pLDDT<70 and high MSA coverage. In total, we assessed 450 variants (**Figure 4A**). Out of these 450 variants, 92 (20.4%) are synonymous, and 27 variants (6%) lead to premature stop codons. The remaining 331 variants (331/450, 73.6%) were classified based on our variant classification scheme. 169 variants (169/331, 51.1%) were predicted to have no effect, 90 (90/331, 27.2%) variants were classified as causing compromised DNA binding, and 67 variants (67/331, 20.2%) were classified as causing unfolding of the domain. For 5 variants (1.5%), we provide no prediction (**Figure 4B**; **Supplemental Table 4**).

**Figure 4.**
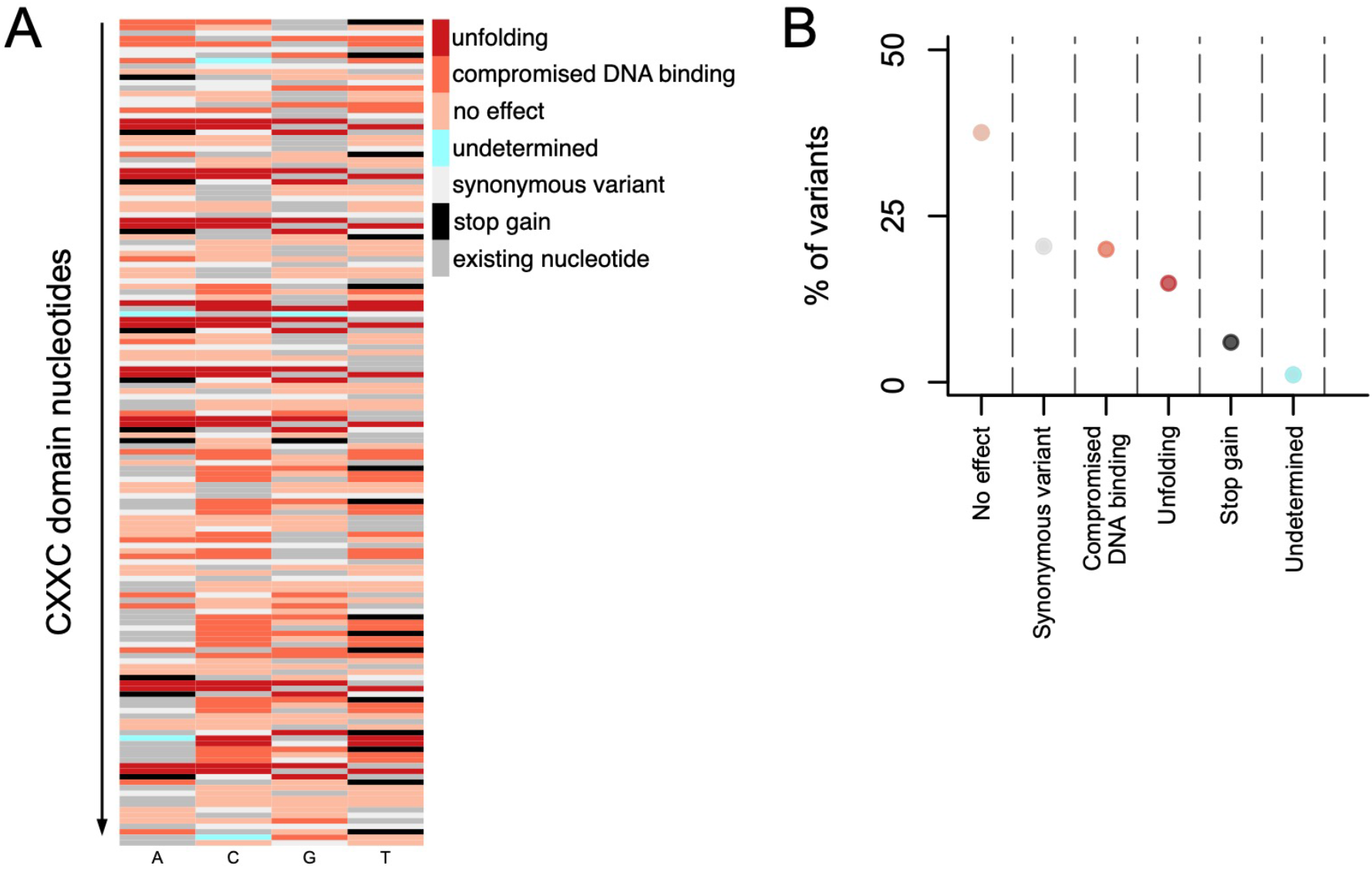
An *in silico* saturation mutagenesis of the CXXC domain of KMT2A. **(A)** Heatmap depicting the predicted effect for each variant. **(B)** The percentage of variants with different effects.

## Discussion

Our contribution in this work is twofold. First, we provide strong genetic evidence that the domains most important for mediating the causal role of KMT2A in Wiedemann-Steiner syndrome are the CXXC domain and – to a lesser extent – the fourth and first PHD fingers. It is noteworthy that these PHD fingers have been shown to be important for stabilizing the interaction between the N- and the C-terminal KMT2A fragments [7]; their enrichment for WSS-causing missense variants may thus not be attributable to their histone-binding function. We emphasize here that our domain fold-enrichment estimates are based on a relatively low number of WSS variants and may change once more variants are reported. However, we anticipate the rank-ordering of the different domains, and the finding that the CXXC is by far the most enriched domain, to remain unchanged.

Our ChIP- and RNA-seq results suggest that lack of KMT2A recruitment to CpG-rich locations – either by missense variants in the CXXC domain or by loss-of-function variants – is central to WSS pathogenesis. However, they also suggest that the phenotype is not mediated via ensuing defects in the deposition of histone methylation, but rather that alternative mechanisms might be at play. This is consistent with the lack of overrepresentation of WSS missense variants within the enzymatic SET domain of KMT2A, and stands in contrast to what was previously observed in Kabuki syndrome, a Mendelian disorder of the epigenetic machinery with considerable phenotypic overlap with WSS caused by variants in KMT2D, which is structurally similar to KMT2A [8].

Our second contribution is the demonstration that the recent breakthrough in protein structure prediction by AlphaFold2 can be leveraged in order to classify the effects of missense variants with experimental-level accuracy. We highlight that here we use AF2 to directly predict the structure of the mutant proteins; based on these mutant structures, we then assess the effect of variants on secondary structural features and electrostatic potential. This is in contrast to recent work that only uses the wild-type structures as input to algorithms that predict the effect of variants on biophysical attributes like ΔΔG [9]. Prior to our study, it has been unclear if AlphaFold2 is capable of predicting mutant structures accurately. While we do not directly assess this, our results indirectly suggest that it may indeed be true. This conclusion is supported by the high concordance between our predictions and relatively extensive experimentally validated effects, as well as by the fact that nearly all variants present in healthy population databases are classified as benign.

In summary, our work yields insights into the pathogenesis of Wiedemann-Steiner syndrome and presents a strategy for characterizing variant effects using AlphaFold2 that we anticipate will be broadly applicable to many other disease-relevant proteins.

## Materials and Methods

### Missense Variants

Missense variants (MVs) present in the general population were obtained from the Genome Aggregation Database (gnomAD; version 2.1.1 and 3.1.1) [10] and the Trans-Omics for Precision Medicine database (TOPmed) [11]. To ensure that these MVs are not deleterious, we only retained those with frequency greater than 0.0001. Somatic MVs were obtained from the Catalogue of Somatic Mutations in Cancer (COSMIC; version 94) [12]. MVs present in individuals with disease phenotypes (Wiedemann-Steiner syndrome, Hereditary sensory neuropathy type 1E, Childhood-onset dystonia, and Beck-Fahrner syndrome) were obtained from ClinVar [13]. To increase our power, we chose to include variants labeled as Pathogenic, Likely Pathogenic, or Variants of Uncertain Significance (VUS), and filtered them using the phred-like CADD scores, acquired from Ensembl Variant Effect Predictor (VEP) [14]. Specifically, we only retained MVs with a phred-like CADD score above 20 [15]. In the case of WSS, we also obtained 16 additional MVs from Lebrun et al, Baer et al, Miyake et al, WD Jones **(Supplemental Table 1)** [16-19]. Disease MVs that were also present in gnomAD were excluded from subsequent analyses. To ensure that our domain enrichment estimates are not artifacts driven by the inclusion of VUS’s, we also performed the domain enrichment analysis after excluding VUS’s and obtained very similar results, with an even greater enrichment in the CXXX domain **(Supplemental Figure 1D)**.

For the enrichment analysis in the CXXC domain of *KMT2B* and *TET3* (**Figure 1C**), we tested if the observed lack of enrichment can be attributed to the low number of total counts in these genes (22 and 16, respectively). In both cases, we observe 0 MVs in the CXXC domain. In contrast, out of the MVs in gnomAD, 0.7% and 2.5% fall within the CXXC domain of *KMT2B* and *TET3*, respectively. Using these gnomAD percentages and the well-known formula for the probability mass function of the binomial distribution, we calculated that, even if the true ratio of the percentage of disease variants falling in the CXXC domain to the corresponding percentage of gnomAD variants is 3 times less compared to the ratio for *KMT2A*, the probability of observing at least one MV in the CXXC domain is 84.5% for *KMT2B* and 99.9% for *TET3*. These estimates make inadequate power an unlikely explanation for the lack of MV enrichment in the CXXC domain of *KMT2B* and *TET3*.

For **Figure 1A** and **Supplemental Figure 2A**, variants in *KMT2A* were plotted using the Mutation Mapper tool from cBioPortal [20].

### ChIP-seq analysis

Raw ChIPseq data (fastq files) were downloaded from GSE99250 [4]. Reads were aligned to the mouse mm10 (GRCm38) reference genome with Bowtie2 using the default settings with the -U option for aligning unpaired reads, generating a sam file output for each fastq file [21]. The sam files were converted to bam files using Samtools view with the -b option, then sorted using Samtools sort with the -O BAM option for outputting bam files [22]. Peaks were called using MACS2, using default settings with a threshold of q-value (-q option) 0.1 for significant peaks [23]. The sorted bam files and the lists of significant peaks (xls files from MACS2) were used as input to the DiffBind package in R [24]. DiffBind was then run with default settings. Differential peaks obtained in DiffBind were given genomic annotations and converted to GRanges objects using the ChIPseeker package in R [25]. Sequences of peaks were obtained using the getSeq() function from the BSgenome.Mmusculus.UCSC.mm10 package in R [26]. The observed-to-expected CpG ratio was calculated using the following formula [27]:

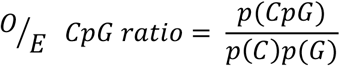

where *p*(*CpG*) represents the proportion of CpGs in a given region (similarly for *p*(*C*) and *p*(*G*)).

### RNA-seq analysis

Raw RNA-seq data (fastq files) were downloaded from GSE99250 [4]. Reads were pseudo-aligned to the mouse mm10 (GRCm38) reference transcriptome using Kallisto [28] (kallisto quant command) with the options --single for single-end reads, -l 150 as an approximation of fragment length, -s 20 as an approximation of fragment length standard deviation, and -b 100 for running 100 bootstraps. The Kallisto output was imported into R using the tximport package [29], with transcripts mapped to genes using the BiomaRt package in R [30]. The tximort software was run with options, type = “Kallisto” and ignoreTxVersion =TRUE. The data was filtered to exclude genes with less than 10 counts. Differential expression analysis was performed using DESeq2 with default settings [31].

For **Figure 2A** and **Figure 2C**, the percentage of differentially marked H3K4me1 peaks and differentially expressed genes were estimated from the corresponding p-value distributions using Storey’s method [32, 33], as implemented in the qvalue R package. Specifically, we used the pi0est() function, with the “pi0.method” parameter set to “bootstrap”.

### Variant effect prediction

We obtained experimentally determined effects of MVs in the CXXC domain of KMT2A from Allen et al and Cierpicki et al [5, 6]. In the absence of experimental data, we chose to not provide a prediction for variants that cause substitutions of salt bridge residues to other amino acids that have been implicated in salt bridge formation (Asp, Glu, Lys, Arg, His) [34]. We classified variants causing substitutions of residues responsible for forming direct hydrogen bonds with the DNA to amino acids that are incapable of forming hydrogen bonds (Ala, Cys, Gly, Ile, Leu, Met, Phe, Pro, Val) as resulting in compromised DNA binding while other substitutions at these residues are applied to the scheme [35].

Structure predictions from AF2 were generated using AF2 Colaboratory (v2.0) [36]. We do not provide a prediction for variants which result in a structure where: a) the pLDDT value of the beta sheets, positioned at the distal ends, drops below 70, or b) the pLDDT value drops to 85 or lower for more than two residues that are not positioned and the distal ends of the domain, or at residues that fall within a secondary structure that is predicted to be absent. Moreover, we do not provide a prediction for the single variant (R1151P) which causes major steric hindrance in the DNA binding face of the domain, since we do not have data to determine the potential impact of this variant.

Secondary structure visualization and computation of mean Coulombic values were performed using UCSF ChimeraX (version 1.2.5) [37], with the AF2 predicted structure as input. MVs were classified as causing a change in electrostatic potential if the mean electrostatic potential value of the mutant structure deviated by more than 0.2 (in either direction) from the wild-type domain, whose mean value is equal to 6.28. We note that the majority of MVs that were classified as disrupting the electrostatic potential caused changes approximately equal to 0.5, with the minimum being 0.31. Finally, functionally important residues were defined as the residues that have been experimentally determined to be implicated in electrostatic interaction with the DNA backbone, residues responsible for hydrogen bond formation with the DNA and residues at the structurally important KFGG site (**Supplemental Table 2**) [6].

## Acknowledgements

TR is supported by a grant from the Wiedemann-Steiner Foundation. HTB is supported by grants from the Louma G. Foundation, the Icelandic Research Fund (#217988, #195835, #206806) and the Icelandic Technology Development Fund (#2010588). Research reported in this publication was supported by the National Institute of General Medical Sciences of the National Institutes of Health under award number R01GM121459.

