## Supplemental Figures for "Missense variants causing Wiedemann-Steiner syndrome preferentially occur in the KMT2A-CXXC domain and are accurately classified using AlphaFold2"

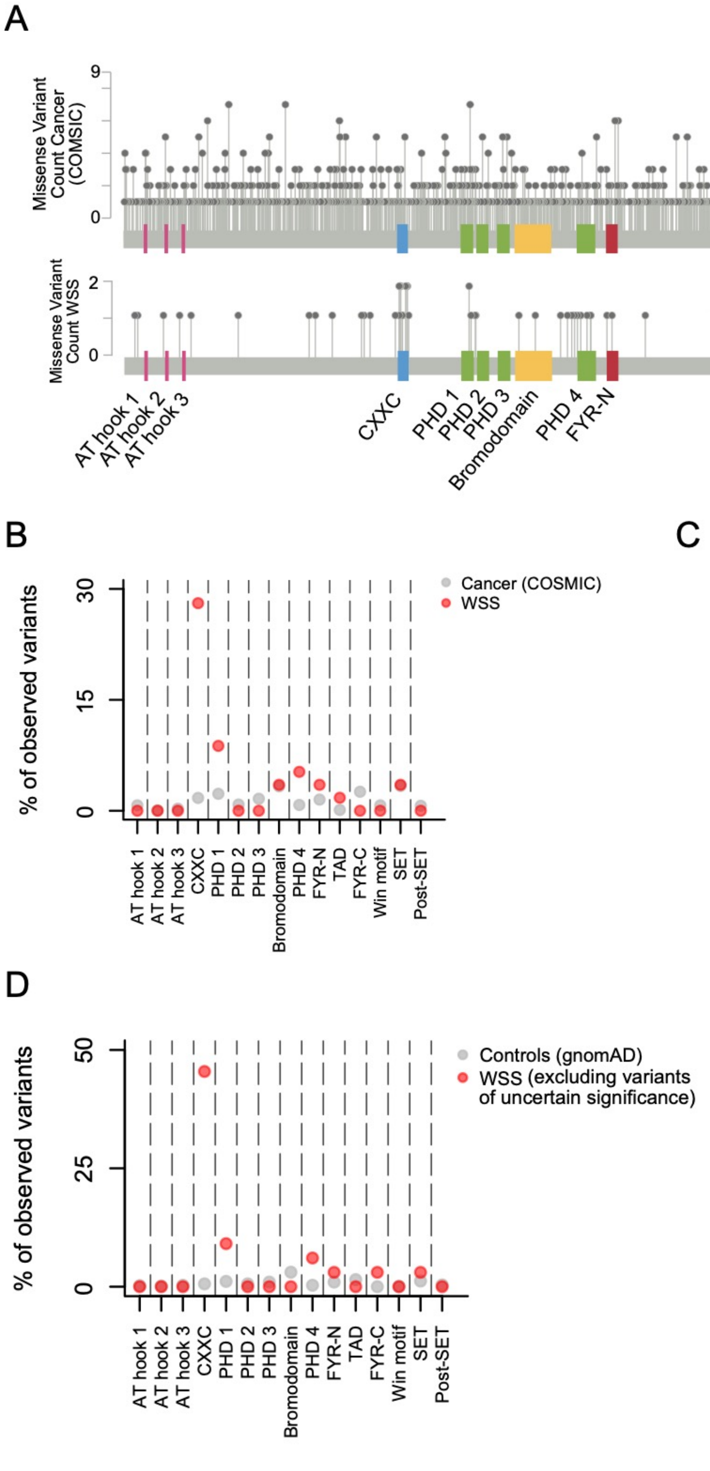

A
Missense Variant Count Cancer (COSMIC)
9
0
Missense Variant Count WSS
2
0
AT hook 1
AT hook 2
AT hook 3
CXXC
PHD 1
PHD 2
PHD 3
Bromodomain
PHD 4
FYR-N
TAD
FYR-C
Win motif
SET
Post-SET
B
% of observed variants
30
15
0
Cancer (COSMIC)
WSS
C
Fold-enrichment (WSS vs COSMIC)
Fold-enrichment (WSS vs gnomAD)
60
30
0
CXXC
PHD 4
D
% of observed variants
50
25
0
Controls (gnomAD)
WSS (excluding variants of uncertain significance)

**Supplemental Figure 1. (A)** KMT2A missense variants in COSMIC (top) and WSS (bottom). **(B)** The percentage of missense variants from COSMIC (grey dots) and WSS (red dots) that fall in each of the different domains of KMT2A. **(C)** Fold-enrichment (odds ratios) of each domain of KMT2A for WSS missense variants, when compared against missense variants from gnomAD (x axis) or COSMIS (y axis). **(D)** The percentage of missense variants from gnomAD (grey dots) and WSS (red dots) that fall in each of the different domains of KMT2A, after excluding variants of uncertain significance from the WSS variants. [CXXC domain odds ratio = 135.4, Fisher's exact test, $p < 2.2e-16$; PHD finger 1 odds ratio = 8.8, Fisher's exact test, $p = 0.008076$; PHD finger 4 odds ratio = 21.3, Fisher's exact test, $p = 0.007914$].

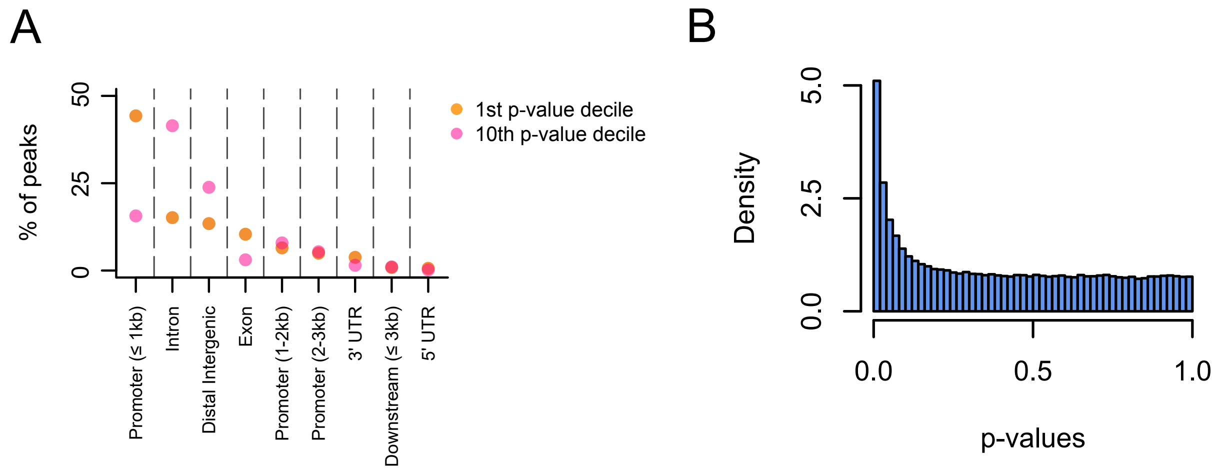


**Supplemental Figure 2. (A)** Genomic annotation of peaks within the 1st (orange dots) and 10th p-value decile (pink dots) from the H3K4me1 differential analysis**. (B)** Histogram of the p-values from the H3K4me1 differential analysis.

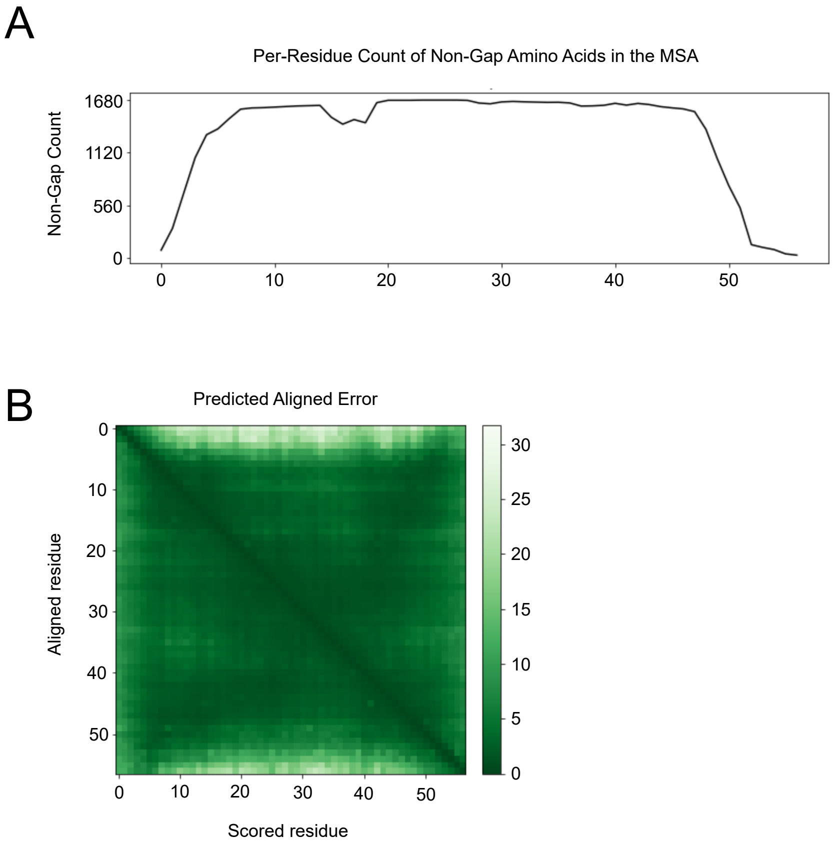


**Supplemental Figure 3. (A)** Multiple sequence alignment depth plot and **(B)** predicted alignment error of KMT2A-CXXC wild-type prediction from AlphaFold2.

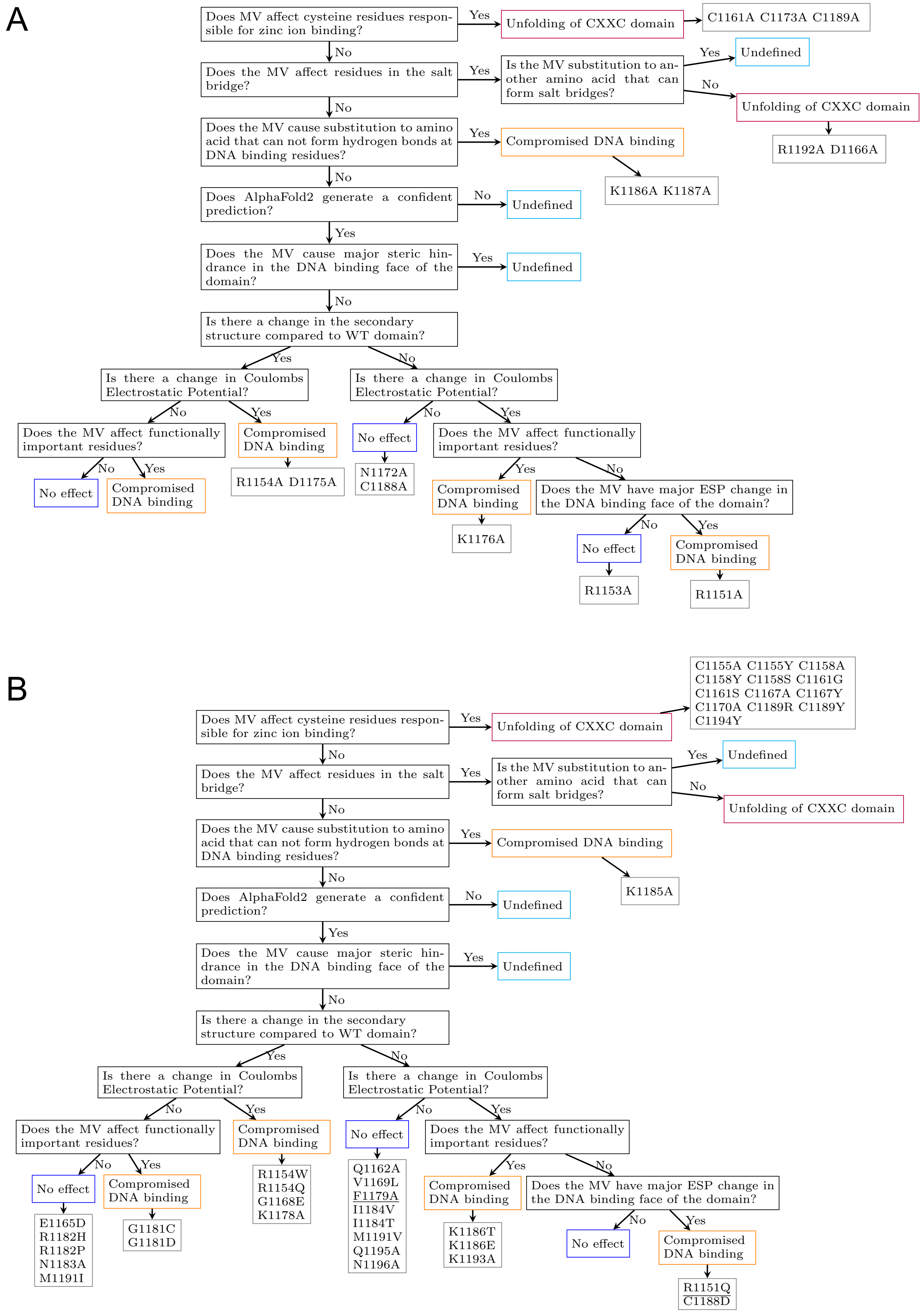
A
Does MV affect cysteine residues responsible for zinc ion binding?
Yes
Unfolding of CXXC domain
C1161A C1173A C1189A
No
Does the MV affect residues in the salt bridge?
Yes
Is the MV substitution to another amino acid that can form salt bridges?
Yes
Undefined
No
Unfolding of CXXC domain
R1192A D1166A
No
Does the MV cause substitution to amino acid that can not form hydrogen bonds at DNA binding residues?
Yes
Compromised DNA binding
K1186A K1187A
No
Does AlphaFold2 generate a confident prediction?
No
Undefined
Yes
Does the MV cause major steric hindrance in the DNA binding face of the domain?
Yes
Undefined
No
Is there a change in the secondary structure compared to WT domain?
Yes
No
Is there a change in Coulombs Electrostatic Potential?
Is there a change in Coulombs Electrostatic Potential?
No
Yes
No
Yes
Does the MV affect functionally important residues?
Compromised DNA binding
R1154A D1175A
No effect
N1172A C1188A
Does the MV affect functionally important residues?
No
Yes
No effect
Compromised DNA binding
Yes
No
Compromised DNA binding
K1176A
Does the MV have major ESP change in the DNA binding face of the domain?
No
Yes
No effect
R1153A
Compromised DNA binding
R1151A
B
Does MV affect cysteine residues responsible for zinc ion binding?
Yes
Unfolding of CXXC domain
C1155A C1155Y C1158A C1158Y C1158S C1161G C1161S C1167A C1167Y C1170A C1189R C1189Y C1194Y
No
Does the MV affect residues in the salt bridge?
Yes
Is the MV substitution to another amino acid that can form salt bridges?
Yes
Undefined
No
Unfolding of CXXC domain
No
Does the MV cause substitution to amino acid that can not form hydrogen bonds at DNA binding residues?
Yes
Compromised DNA binding
K1185A
No
Does AlphaFold2 generate a confident prediction?
No
Undefined
Yes
Does the MV cause major steric hindrance in the DNA binding face of the domain?
Yes
Undefined
No
Is there a change in the secondary structure compared to WT domain?
Yes
No
Is there a change in Coulombs Electrostatic Potential?
Is there a change in Coulombs Electrostatic Potential?
No
Yes
No
Yes
Does the MV affect functionally important residues?
Compromised DNA binding
R1154W R1154Q G1168E K1178A
No effect
Q1162A V1169L F1179A I1184V I1184T M1191V Q1195A N1196A
Does the MV affect functionally important residues?
No
Yes
No effect
E1165D R1182H R1182P N1183A M1191I
Compromised DNA binding
G1181C G1181D
Yes
No
Compromised DNA binding
K1186T K1186E K1193A
Does the MV have major ESP change in the DNA binding face of the domain?
No
Yes
No effect
Compromised DNA binding
R1151Q C1188D

**Supplemental Figure 4.** The classification of individual missense variants in the **(A)** training set and **(B)** test set. Variants in the test set that are classified incorrectly are underlined.
