## Supplemental Table 2 for "Missense variants causing Wiedemann-Steiner syndrome preferentially occur in the KMT2A-CXXC domain and are accurately classified using AlphaFold2"

Functionally important residues in the CXXC domain of KMT2A

| | |
|---|---|
| Residues implicated in electrostatic interaction with the DNA | Arg1150, Arg1154, Lys1176, Lys1178, Lys1185, Lys1190, Arg1192, Lys1193 |
| Residues responsible for hydrogen bond formation with the DNA | Lys1185, Lys1186, Gln1187 |
| Residues at the KFGG site | Lys1178, Phe1179, Gly1180, Gly1181 |
